# Real-time precision opto-control of chemical processes in live cells

**DOI:** 10.1101/2022.01.23.477373

**Authors:** Matthew G. Clark, Gil Gonzalez, Jesus Aldana-Mendoza, Mark S. Carlsen, Gregory Eakins, Chi Zhang

## Abstract

Precision control of molecular activities and chemical reactions in live cells is a long-sought capability by life scientists. No existing technology can probe molecular targets in cells and simultaneously control the activities of only these targets at high spatial precision and on the fly. We develop a real-time precision opto-control (RPOC) technology that detects a chemical-specific optical response from molecular targets during laser scanning and uses the optical signal to trigger an acousto-optic modulator, which allows a separate laser beam to only interact with the molecules of interest without affecting other parts of the sample. RPOC can automatically probe and control biomolecular activities and chemical processes in dynamic living samples with submicron spatial accuracy, nanoseconds response time, and high chemical specificity.

## Introduction

The advancement of microscopy technologies has revealed unprecedented details of biological processes with superb resolution and chemical information. However, the capability to control chemical processes in live cells with high spatial accuracy and molecular selectivity in real-time is still lacking. Conventional chemical treatment by culturing cells with chemical compounds has poor spatial delivery selectivity and might pose off-target effects for the accurate understanding of compound-target interactions. Genetic methods such as CRISPR and RNA interference can control the expression and activity of proteins *(1-3)*. However, transfection and incubation require sophisticated pre-preparation and passaging processes with little temporal and spatial control. Optical tweezers and trapping can only physically manipulate a few pre-detected targets (*4-6*). Current laser ablation and manipulation methods are based on pre-image acquisition and manual operation of laser beams to interact with the target-of-interest (*7-10*). Optogenetics methods can control functions of neurons using light radiation and light-sensitive ion channels, however, require pre-imaging and demonstrate little sub-cellular precision (*11-13*). Thus, the existing optical manipulation technologies cannot apply to highly dynamic living biological samples to control molecular activities with high spatial accuracy and chemical specificity.

In this work, we develop a real-time precision opto-control (RPOC) technology that can detect and control molecules simultaneously, selectively, and precisely at the only desired activity sites. First, during laser scanning, an optical signal is generated at a specific pixel from target molecules. Then, the detected optical signal is compared with preset values using comparator circuitry. A desired optical signal will activate an acousto-optic modulator (AOM) which is used as a fast switch to couple another laser beam to interact at the same pixel. The optical signal detection, processing, and opto-control happen within 30 ns and in real-time during laser scanning. Digital logic functions allow opto-control of molecular activities based on the logic output from multiple signal channels. RPOC can accurately detect and control biomolecules in real-time without affecting other locations in the system. It is highly chemically selective since the optical signal can be selected from a range of responses such as fluorescence and Raman. This technology offers an unprecedented way to automatically and selectively control molecular activities and chemical reactions with sub-micron spatial precision.

## Results

### The RPOC platform

The concept of RPOC, which is based on fast laser scanning, is illustrated in **Fig. 1, A and B**. A laser(s) for optical signal excitation is scanning through the field of view. During the laser scanning, if a chemical-specific optical signal is detected and satisfies a preset condition (e.g. surpasses a threshold value), it will trigger an AOM to send a separate laser beam to interact with the sample at the same pixel in real-time. Optical signals that do not satisfy this condition will “turn off” the control laser beam to avoid laser interaction. Digital comparator circuits (**fig. S1**) were designed for presetting the selection conditions (e.g. the threshold V_T_), performing analog/digital comparisons, and sending out a standard transistor-transistor logic (TTL) voltage for AOM control. A schematic of the RPOC system is shown in **Fig. 1C**. A dual-output femtosecond laser is used to perform optical signal excitation and opto-control. The 1045 nm laser output is used as the Stokes beam and the frequency-tunable laser output is used as the pump beam for stimulated Raman scattering (SRS) signal generation (*14, 15*). The laser beams are also used for two-photon excitation fluorescence (TPEF) signal excitation (*16*). Portions of both outputs are frequency-doubled to the visible range for opto-control. The selected control laser is sent to an AOM and commanded by the comparator circuits. The laser beam profiles after the AOM using ‘0’ and ‘1’ TTL commands are shown in **Fig. 1, D and E**, respectively. The 0^th^ order output is blocked by a beam stop and the 1^st^ order of the AOM output is combined with the excitation IR laser beams by a dichroic mirror before coupling into the microscope. **Fig. 1F** shows the ∼15 ns response time of the comparator circuits. The AOM response time (**Supplementary material**) is calculated to be ∼7 ns. Therefore, the response time of the opto-control system is less than 30 ns, much shorter than the 10 µs pixel dwell time for laser scanning. The spatial resolution of the SRS and TPEF modalities is measured to be 373 nm (**fig. S2**). The PROC control laser beam gives a spatial precision of 525 nm (**fig. S3**).

**Fig. 1.**
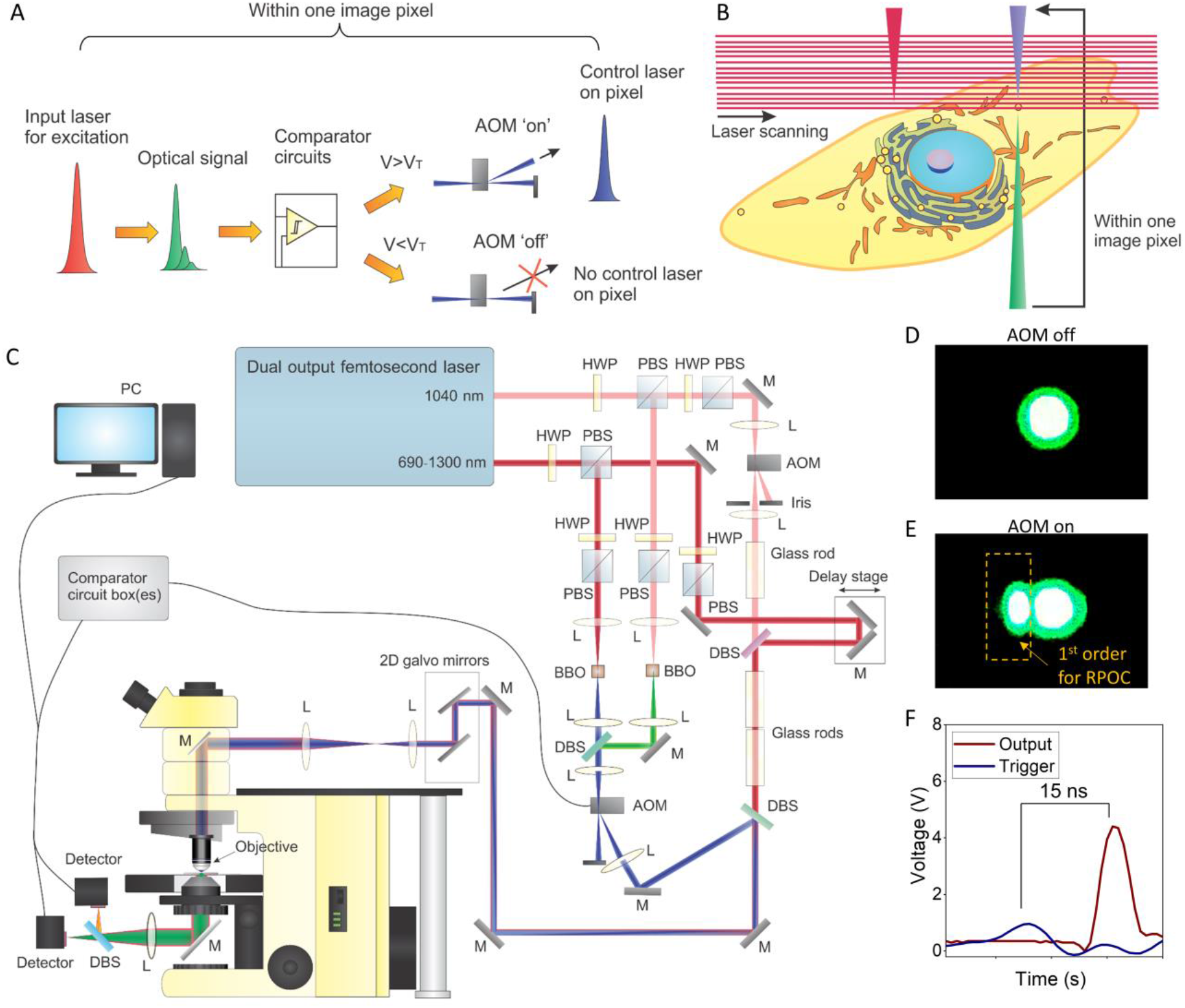
The RPOC concept and optical configuration. (A) An illustration of the RPOC concept. (B) The illustration of RPOC for selective control of molecular activities in a cell during laser scanning. (C) A schematic of the RPOC experimental setup. HWP, half-wave plate; L, lens; M, mirror; DBS, dichroic beam splitter; PBS, polarization beam splitter; AOM, acousto-optic modulator. (D) The profile of the control laser beam (at 522 nm) after the AOM when the AOM is turned off. (E) The profile of the control laser beam after the AOM when the AOM is turned on (the 1^st^ order deflection is highlighted). (F) The response time of the comparator box is measured to be ∼15 ns.

### Real-time control of active pixels (APXs) using chemical-specific optical signals

An ‘active pixel’ (APX) is defined as the pixel location at which the control laser beam is turned on. Tracking APXs is critical for visualizing the opto-control locations. In the over-sampling condition (the pixel size is smaller than the laser beam size at the focus), the size of the laser interaction area is larger than the size of the APXs (**Fig. 2A**). Higher intensity thresholds reduce APXs and laser interacting areas. Similarly, at the same intensity threshold, weaker optical signals above the threshold reduce APXs and the laser interaction areas (**Fig. 2B**). Increasing the pixel size might change these properties. Different optical signal intensities that would result in different APXs in the oversampling condition might give the same APXs and interaction areas (**Fig. 2C**). When the pixel size is greater than the actual laser beam size, the APXs might be the same or even larger than the actual interaction area (**Fig. 2C**). The oversampling condition is used throughout this work.

**Fig. 2.**
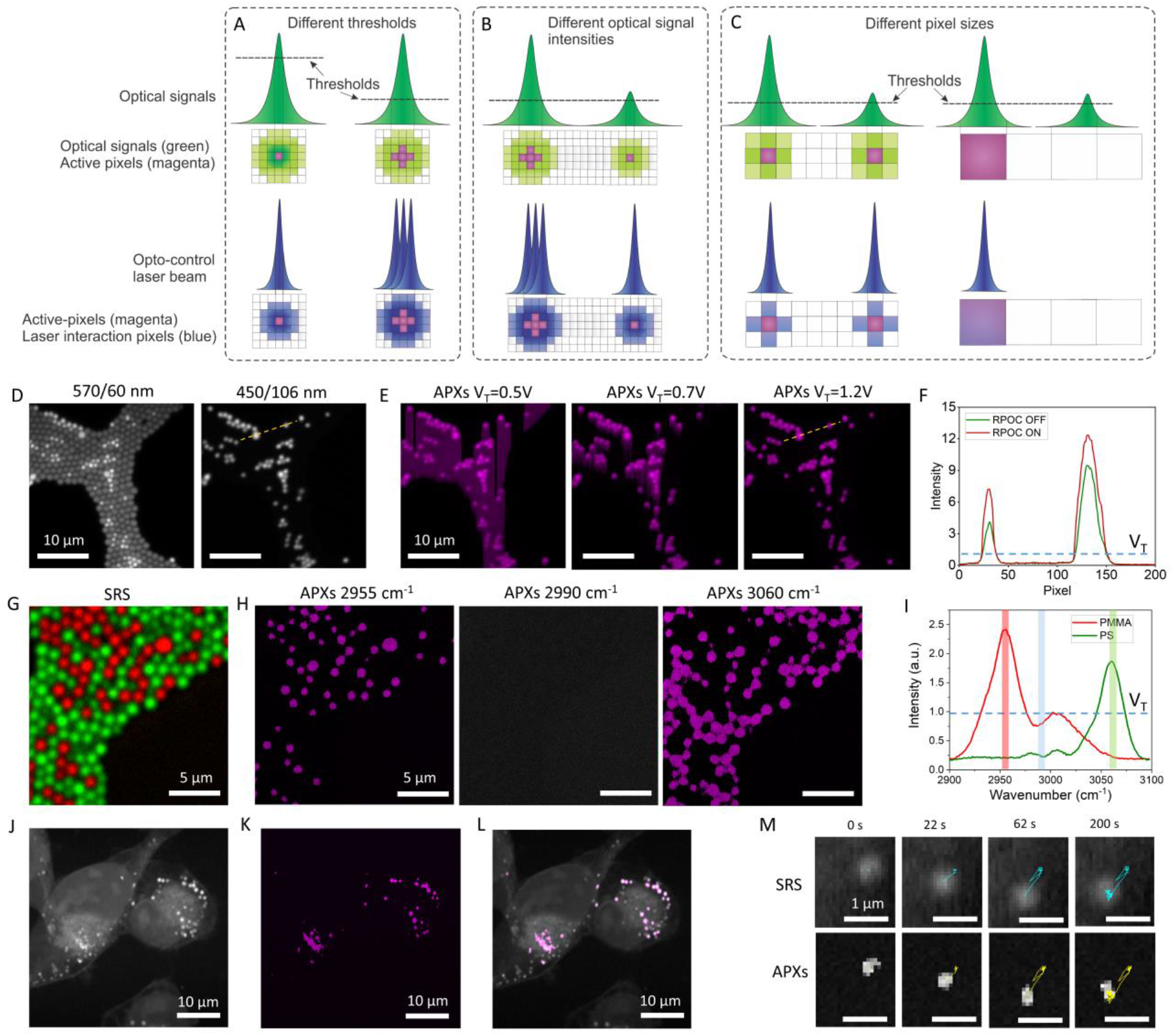
Mapping out RPOC APXs. (A) An illustration of APX selection using different signal thresholds in the over-sampling condition. The green pulses and pixels indicate optical signals, the magenta pixels indicate APXs, and the blue pulses and pixels indicate the interaction pulses and pixels. (B) An illustration of APX selection using different optical intensities in the over-sampling condition. (C) An illustration of APX selection at larger pixel sizes for different optical signals. (D) Mixed fluorescent particles detected in the 570/60 nm channel (left, orange and green fluorescent particles) and 450/106 nm channel (right, only the green fluorescent particles). (E) APXs determined using different V_T_ values for the signals from the 450/106 nm channel. (F) Comparing the 450/106 nm channel optical intensity when the RPOC is turned on (red) and off (green). (G) A pseudo-color SRS image containing PMMA (red) and PS (green) particles. (H) APXs determined using the PS peak at 2950 cm^-1^ (left), PMMA peak at 3060 cm^-1^ (right), and no Raman peaks at 2990 cm^-1^ (middle). (I) SRS spectra of PMMA (red) and PS (green). The red, green, and blue lines are wavenumbers used for RPOC in panel H. (J) An SRS image of MIA PaCa-2 cells in the lipid CH_2_ stretching region. (K) APXs determined using SRS signals from LDs. (L) An overlay of the SRS image and the APXs turned on only at the LDs. (M) Time-lapse SRS images of a LD in a live MIA PaCa-2 cell (top row) and the corresponding APXs determined by the SRS signals (bottom row). The color curves plot trajectories of the LD and the APXs in 200 s.

We first use fluorescence signals from microparticles to determine APXs. Laser pulses at 800 nm is used to excite the TPEF signals of mixed 1 µm particles. **Fig. 2D** shows fluorescence signals detected in the 570/60 nm (left panel) and the 450/106 nm (right panel) fluorescence channels. The latter is paired with a comparator circuit box to determine the APXs. The selection condition is the fluorescence signal above a preset threshold V_T_. When the V_T_ is low (0.5 V and 0.7 V), APXs exceed the fluorescence pixels in the 450/106 nm channel (**Fig. 2E**). At a proper threshold V_T_=1.2 V, APXs perfectly match the fluorescence signals. **Fig. 2F** compares the intensity profiles at selected lines in panels A and B when RPOC is turned off and on. The intensity increase is contributed by the leaking of the control laser which is only turned on by the fluorescence signal at the particles.

**Fig. 2G** shows chemical maps of mixed poly(methyl methacrylate) (PMMA) and polystyrene (PS) particles generated by hyperspectral SRS microscopy (*17*). Using Raman shifts at 2955 cm^-1^ or 3060 cm^-1^, we can determine APXs using either PMMA or PS SRS signals (**Fig. 2H**). The selection condition is optical signals greater than V_T_=1 V. **Fig. 2I** displays the SRS spectra of PMMA and PS, the V_T_, and the selected Raman shifts for APX determination in **Fig. 2E**. APXs can be selected on different chemicals in real-time by tuning laser frequencies to match different Raman transitions (**movie S1**).

Using the lipid CH_2_ symmetric stretching SRS signals at 2855 cm^-1^, we can automatically select APXs only at the lipid droplets (LDs) in live cells, as shown in **Fig. 2, J-L**. Side-by-side time-lapse images of SRS and APX selection using the lipid signals are shown in **movie S2**. The trajectory of APXs triggered by a single LD matches the corresponding LD trajectory (**Fig. 2M, movie S3**). We also demonstrate 3D precision controlling of the APXs in MIA PaCa-2 cells, as shown in **movies S4 and S5**. These results highlight the capability of tracing intercellular dynamics for APX determination using RPOC.

### Digital logic control of APXs

A second comparator box with digital logic functions is also design as shown in **fig. S4**. digital logic functions can be selected from AND, OR, NAND, and NOR. Using the two comparator circuit boxes, we can select any intensity range from a single detector for APX determination. The connections to achieve this function are illustrated in **Fig. 3A** and **fig. S5. Fig. 3B** shows APXs selected on the LDs using only comparator box 1 and a single intensity threshold. **Fig. 3C** shows APXs determined using different intensity passbands between the upper and lower thresholds. The APXs selected between V_T_=0.2-0.3 V are more associated with the endoplasmic reticulum (ER) and between 0.1-0.16 V are mostly on cytosols. Spectral phasor analyses of hyperspectral SRS images of the same cells, which segment different cellular compartments (*18*), are shown in **fig. S6** for comparison.

**Fig. 3.**
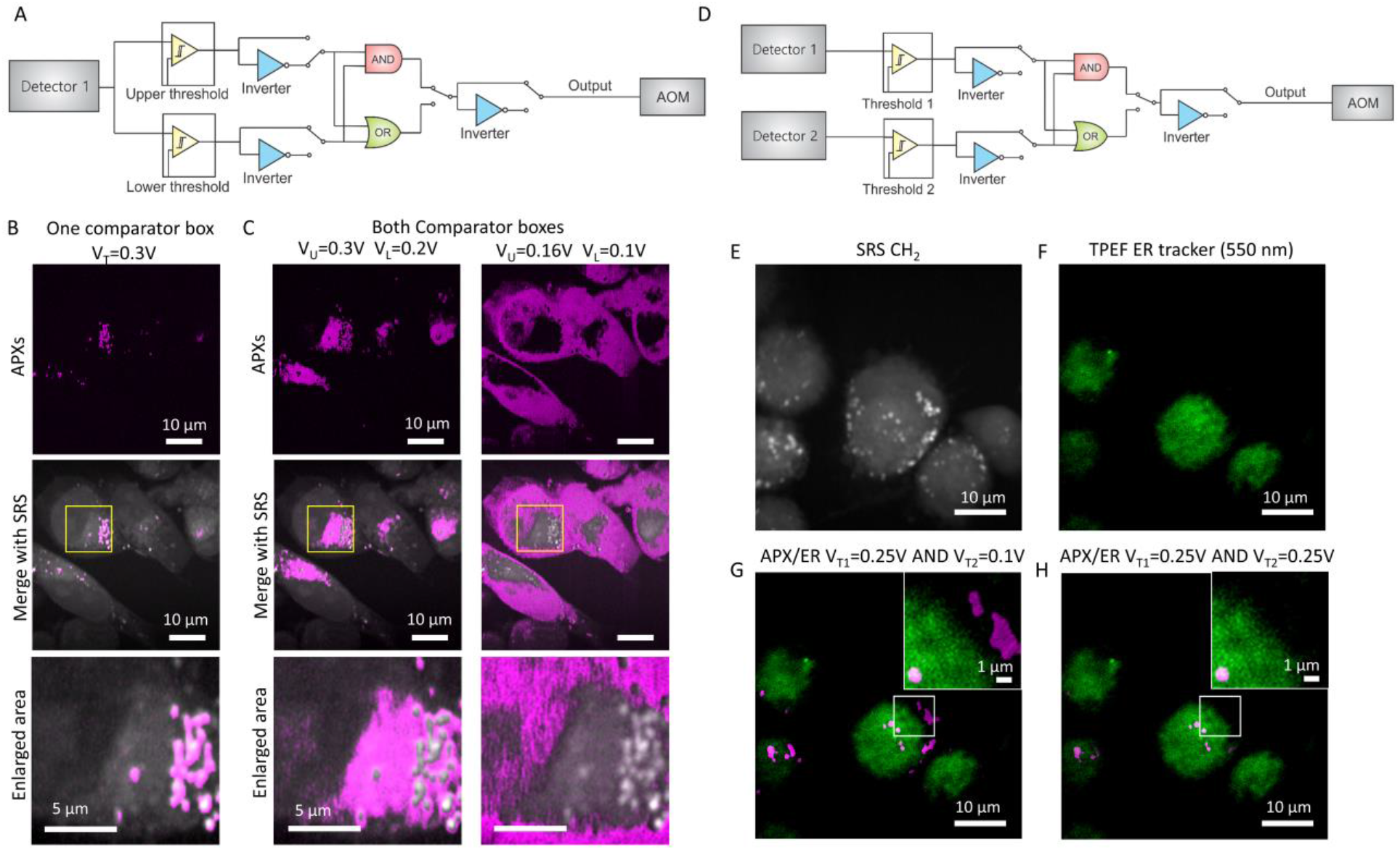
Digital logic control of APXs. (A) Illustration of electronics to select an intensity passband for determination of APXs. (B) The APXs selected using only one comparator box with V_T_=0.3 V (upper panel), the overlay of the APXs with the corresponding SRS image from the same field of view (middle panel), and a magnified image from the selected area (bottom panel). (C) APXs selected using two comparator boxes with the intensity range between 0.2-0.3 V (left panels) and 0.1-0.16 V (right panels). (D) Illustration of electronics to choose the AND function for determination of APXs using two comparator boxes. (E) An SRS image of MIA PaCa-2 cells at the CH_2_ stretching vibration. (F) An TPEF image of the MIA PaCa-2 cells labeled using ER tracker. (G) An overlay of the TPEF image from ER and the APXs determined using V_T1_=0.25 V (SRS) and V_T2_=0.25 V (TPEF). (H) An overlay of the TPEF image from ER and the APXs using V_T1_=0.25 V (SRS) and V_T2_=0.25 V (TPEF).

The connections for implementing the digital logic functions using two comparator boxes and two detectors are illustrated in **fig. S7**. The selection of the AND function is illustrated in **Fig. 3D**. We first demonstrate the AND function using mixed fluorescent PS particles, non-fluorescent PS particles, and nicotinamide adenine dinucleotide hydrogen (NADH) crystals, as shown in **fig. S8 and S9**. The AND function allows determining APXs only on the fluorescent PS particles that show up in both TPEF and SRS channels. Next, we used SRS to excite lipid signals (**Fig. 3E**) in MIA PaCa-2 cells and label the cells using a fluorescent ER tracker which can be visualized in the TPEF channel (**Fig. 3F**). By using an appropriate V_T1_ in the SRS channel and a low V_T2_ in the TPEF channel, APXs can be selected from most of the LDs in the cells (**Fig. 3G**). Increasing the V_T2_ in the TPEF channel can exclude the LDs outside the ER and excite APXs on LDs only on the ER (**Fig. 3H**). These results demonstrate using the AND logic from two separate detectors for APX determination. The connections of OR, NAND, and NOR functions for RPOC are illustrated in **fig. S10**.

### Control and quantification of chemical changes at sub-micron precision

To demonstrate precision control of chemical processes using the RPOC, we used a photochromic molecule, *cis*-1,2-dicyano1,2-bis(2,4,5-trimethyl-3-thienyl)ethene (CMTE), which can be changed from its open *cis* isomer (**1a**) to closed isomer (**1b**) by UV light and switched back by visible light at 520 nm (**Fig. 4A**) (*19, 20*). A strong Raman signature peak at 1510 cm^-1^ can be detected for **1b** but not **1a** (*19*). The SRS signal from this peak can be visualized by tuning the pump beam to 902 nm. We found that the combination of the pump and Stokes pulsed lasers for SRS imaging can also transform CMTE to **1b**. A 522 nm laser beam frequency-doubled from the 1045 nm laser output is used for RPOC to convert **1b** to **1a** at selected subcellular locations. The Raman transition at 1510 cm^-1^ is used as the reporter for RPOC-induced **1b** to **1a** conversion during laser scanning (**Fig. 4B**).

**Fig. 4.**
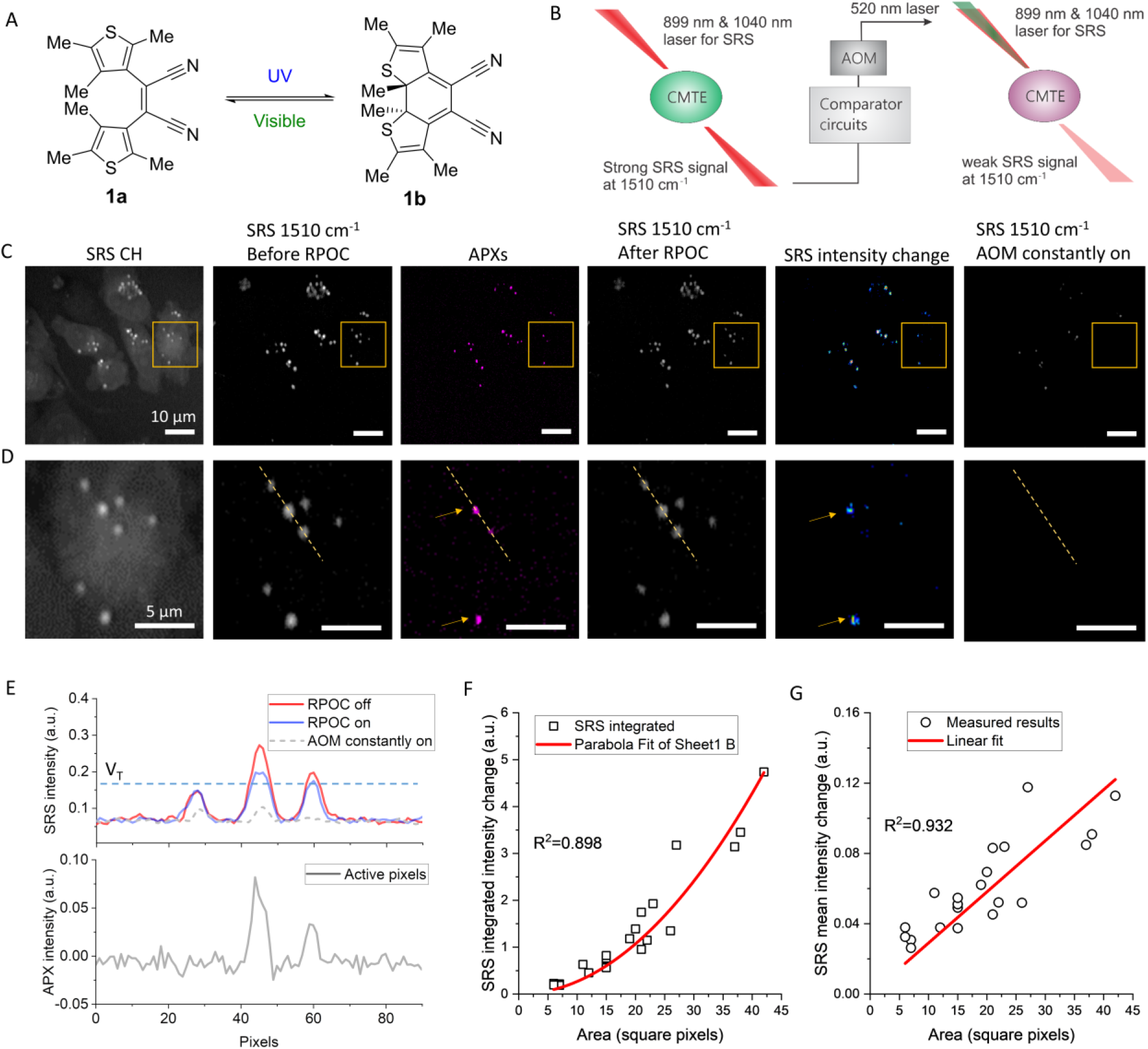
Precision control and quantitative comparison of site-specific chemical changes using RPOC. (A) An illustration of switching states of CMTE between the open *cis* isomer (**1a**) and the closed isomer (**1b**) forms using UV and visible (green) light. (B) An illustration of the workflow for CMTE conversion by the RPOC using 1510 cm^-1^ SRS signal and a 522 nm laser. (C) An SRS image in the CH_2_ region, at 1510 cm^-1^ before RPOC, the APXs, at 1510 cm^-1^ after RPOC using V_T_=0.17 V, the SRS intensity difference before and after RPOC, at 1510 cm^-1^ after the AOM constantly on for 20 frames. (D) Magnified images of the selected areas in panel C. (E) SRS intensity profiles of images and APXs long the dotted lines in panel D. The dashed curve shows the SRS intensity profile along the same line after 20 scan frames with AOM constantly on. (F) Integrated SRS intensity changes of CMTE as a function of the number of APXs for CMTE aggregates. Open circles are experimental results, the curve is the quadratic fitting. (G) Mean SRS intensity changes of CMTE as a function of the number of APXs for CMTE aggregates. Open circles are experimental results, the line is the linear fitting.

First, we treat MIA PaCa-2 cells with CMTE and observed accumulation of CMTE in LDs of the cells due to the hydrophobic structure of the chemical (**Fig. 4, C and D**). Then, we used RPOC to convert **1b** to **1a** at selected locations of the sample with a control laser beam power of ∼10 µW. A single comparator box with a selection condition of V_T_>0.17 V was used to determine APXs which are majorly contributed by high-intensity CMTE aggregates, as shown in **Fig. 4, C and D**. After five frames of RPOC laser scanning, SRS signals at 1510 cm^-1^ from the APX-associated pixels are reduced, which can be visualized from the SRS intensity difference image in **Fig. 4, C and D**. If the control laser beam is constantly turned on during laser scanning for 20 frames, SRS signals at 1510 cm^-1^ on the entire image are significantly reduced (**Fig. 4, C and D**). The pixels where chemical conversion happened, as shown in the SRS intensity difference images, agree with the APXs. **Fig. 4E** plots SRS intensity profiles along the dashed lines in **Fig. 4D**, for images before and after the RPOC, the APXs, and after nonselective laser control after 20 frames. We see that laser-induced chemical changes of CMTE only happen at the APXs. Such chemical changes can be quantified by integrating the SRS intensity change of CMTE on APXs of each aggregate, which shows a quadratic dependence with the number of APXs from each aggregate (**Fig. 4F**). This nonlinear dependence arises from the oversampling condition used for RPOC, as illustrated in **Fig. 2A**. The mean SRS intensity change, on the other hand, has a near-linear dependence with the number of APXs (**Fig. 4G**). These analyses show that RPOC can not only selectively control chemical changes in space but also potentially quantify the amount of products and reaction rates.

### Precision control of molecular activities using different selection conditions

RPOC can control CMTE at different parts of cells using various selection conditions. We first connected two comparator boxes as illustrated in **fig. S3** to select an SRS signal range between two intensity levels. Here, the RPOC is only applied for a single-frame laser scanning with 10 µs dwell time per pixel. **Fig. 5, A and B** display the SRS signals from CMTE at 1510 cm^-1^ before the RPOC, the APXs, after a single frame RPOC, and the SRS intensity changes with a selection condition of V_L_=0.15 V and V_U_=0.4 V, where V_L_ and V_U_ are the lower and upper signal limits for RPOC, respectively. The SRS image of CH_2_ stretching is shown in **fig. S11**. Since V_U_ is very high, this selection range determines the APXs mostly from the centers of the aggregates that contribute to strong CMTE SRS signals (**Fig. 5C**), similar to using a single comparator box as shown in **Fig. 4**. When the selection condition is chosen as V_L_=0.10 V and V_U_=0.12 V, as shown in **Fig. 5, D and E**, weaker SRS signals selected APXs from the edges of most aggregates. In this case, the centers of the aggregates, which are not associated with APXs, are left unchanged, while the molecules at the edges of the aggregates are converted from **1b** to **1a** by RPOC. **Fig. 5F** plots the SRS intensity change and APXs along the dotted line and the intensity thresholds. The CMTE transition from **1b** to **1a** occurs only at the edges of the aggregates. There are pixels with noticeable SRS signal decrease outside APXs (**Fig. 5F**). This is due to the use of an oversampling condition in which the RPOC laser interacting range is larger than the APXs, as illustrated in **Fig. 2A**.

**Fig. 5.**
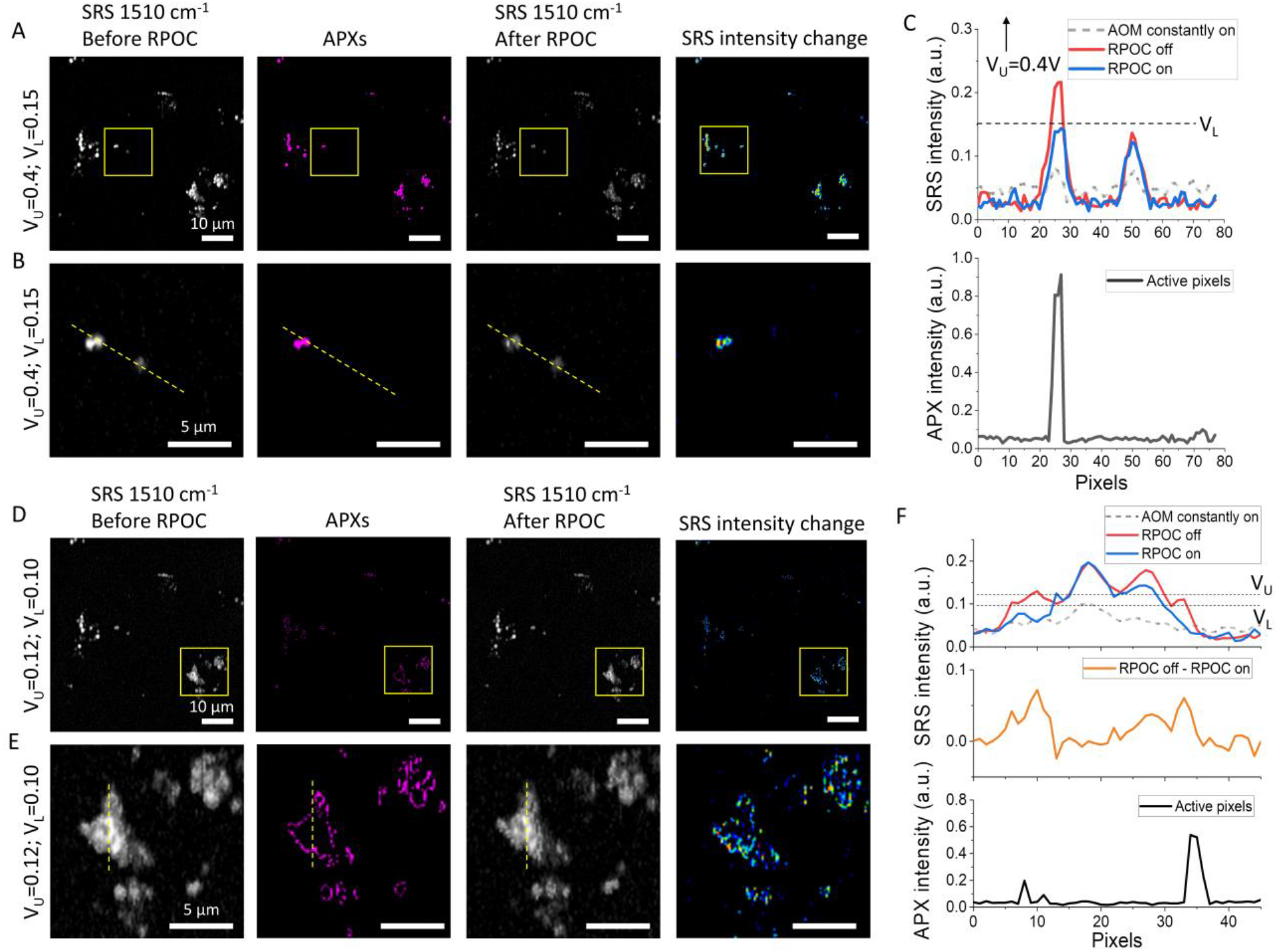
Precision control of chemical changes by RPOC using different optical intensity ranges. (A) An SRS image at 1510 cm^-1^ before RPOC (left), APXs selected by an upper threshold (V_U_) of 0.4 V and lower threshold (V_L_) of 0.15 V (middle left), an SRS image at 1510 cm^-1^ after the RPOC (middle right), and the SRS intensity difference before and after the RPOC (right). (B) Magnified images from the highlighted regions in panel A. (C) SRS intensity profiles of images long the dotted lines in panel B. The dashed curve shows the SRS intensity profile along the same line after 20 scan frames with AOM constantly on. (D) Similar images as in panels A, using V_U_=0.12 V and V_L_=0.10 V for APX selection. (E) Magnified images from the selected regions in panel D. (F) SRS intensity profiles of images long the dotted lines in panel E. The dashed curve shows the SRS intensity profile along the same line after 20 scan frames with AOM constantly on. The middle panel plots SRS intensity difference before and after RPOC.

Next, we used ROPC to selectively control the **1b** to **1a** conversion accumulated only in ER-associated LDs. SRS was used to detect CMTE targeting the 1510 cm^-1^ peak while ER tracker in the TPEF (550-600 nm) channel was used to delineate ER in live MIA PaCa-2 cells. As shown in **Fig. 6, A and B**, APXs can be determined on ER-associated LDs using SRS and TPEF signals from two detectors with the digital AND function and appropriate threshold levels. The ER boundary is shown in **Fig. 6B**. By plotting the SRS intensity profiles of three LDs (**Fig. 6C**) and their corresponding APXs (**Fig. 6D**) along the dashed line (**Fig. 6B**), we find that LD #3, which is not on the ER, is not affected by RPOC; while LDs #1 and #2, both on the ER, are converted from **1b** to **1a** by RPOC. These results demonstrate that digital logic RPOC can control molecular activities and chemical reactions associated with multiple organelles or related to organelle interactions.

**Fig. 6.**
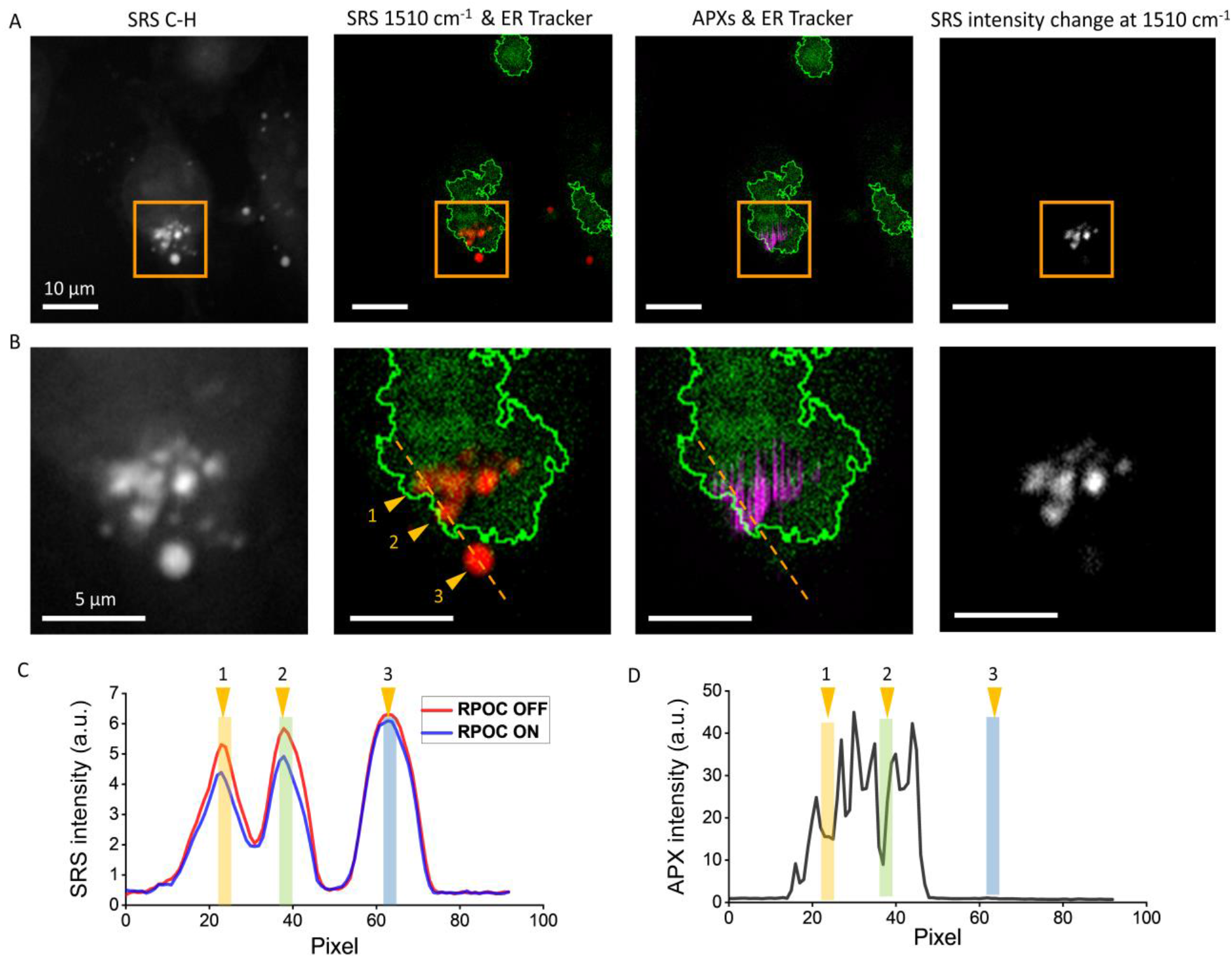
Precision control of chemical changes by RPOC using digital logic from two detectors. (A) Images of MIA PaCa-2 cells showing SRS signals at 2855 cm^-1^ CH stretching, the overlay of SRS signals from CMTE at 1510 cm^-1^ and ER tracker fluorescence signals, the overlay of APXs and ER tracker signals, and the CMTE SRS signal changes after RPOC. (B) Magnified images from the highlighted areas in panel A. (C) CMTE SRS intensity profiles before and after RPOC along the selected lines in panel B. (D) Intensity profiles of APXs. Colored bars highlight positions of selected LDs in panel B.

## Discussion

We for the first time demonstrate real-time precision opto-control of molecular activities and chemical processes triggered by optical signals from the molecules at submicron spatial precision. RPOC can perform active control of light-sensitive molecules and chemical reactions in living biological samples due to the fast response and automatic APX determination. In this work, we majorly focused on demonstrating the RPOC capability using a photoswitchable molecule CMTE. RPOC can also be applied to control newly developed photochromic vibrational probes (*21, 22*), widely used photoswitchable fluorescent molecules (*23, 24*), and light-sensitive chemical reactions (*25-28*) at high spatial and temporal accuracy.

The continuous improvement of RPOC will lead to more opportunities in biophotonics and biological sciences. For example, further optimization of the control laser beam can improve the RPOC precision. Instead of using an expensive femtosecond laser, a more cost-effective and compact RPOC platform can be developed based on continuous-wave (CW) lasers. The CW-RPOC system would greatly reduce the system cost and ideal for integration with commercial fluorescence microscopes. Programmable acousto-optic tunable filters would also allow for the selection of different laser beams automatically for RPOC. Improvement in optics and electronics, such as using an electro-optic modulator and resonant mirrors would further improve the RPOC response time for high-speed laser scanning systems.

PROC offers a way for biologists and chemists to control biomolecular behaviors and chemical reactions precisely and automatically in space and time without affecting unwanted targets. We believe RPOC will have important applications, when combined with photoactivatable molecules, for better control of enzyme activities, high accuracy-controlled release, high precision optogenetics, and improved precision treatment. Applying digital logic functions in RPOC with photoswitchable fluorescent molecules would also enable recording and saving organelle interactions for live systems. Future research will focus on demonstrating the capabilities of RPOC in these applications.

## Author contributions

C.Z. designed the project and experiment. C.Z. and M.G.C. performed the experiments, analyzed the results, and wrote the paper. M.G.C. constructed the optical system for RPOC. G.A.G. helped prepare the cancer cells for imaging. J.A.M helped in biological sample preparation. M.S.C. designed and fabricated the comparator circuits. G.E. designed and fabricated the tuned amplifier and photodetector for SRS signal detection.

## Data and materials availability

All data, code, and materials used in the analysis are available in the corresponding authors’ lab. All data are available in the main text or the supplementary materials.

## References

1. R. Y. Tsien, The green fluorescent protein, Annu. Rev. Biochem., 67, 509 (1998).

2. P. J. Paddison, A. A. Caudy, E. Bernstein, G. J. Hannon, D. S. Conklin, Short hairpin RNAs (shRNAs) induce sequence-specific silencing in mammalian cells, Genes Dev., 16, 948 (2002).

3. R. Barrangou et al., CRISPR provides acquired resistance against viruses in prokaryotes, Science, 315, 1709 (2007).

4. J. Enger, M. Goksör, K. Ramser, P. Hagberg, D. Hanstorp, Optical tweezers applied to a microfluidic system, Lab. Chip, 4, 196 (2004).

5. H. Zhang, K.-K. Liu, Optical tweezers for single cells, J. R. Soc. Interface, 5, 671 (2008).

6. M. Daly, M. Sergides, S. Nic Chormaic, Optical trapping and manipulation of micrometer and submicrometer particles, Laser Photonics Rev., 9, 309 (2015).

7. S. A. Boppart et al., High-Resolution Optical Coherence Tomography-Guided Laser Ablation of Surgical Tissue, J. Surg. Res., 82, 275 (1999).

8. H. He et al., Manipulation of cellular light from green fluorescent protein by a femtosecond laser, Nat. Photonics, 6, 651 (2012).

9. F. Shi et al., Mitochondrial swelling and restorable fragmentation stimulated by femtosecond laser, Biomed. Opt. Express, 6, 4539 (2015).

10. T. Meyer et al., CARS-imaging guidance for fs-laser ablation precision surgery, Analyst, 144, 7310 (2019).

11. K. Deisseroth et al., Next-Generation Optical Technologies for Illuminating Genetically Targeted Brain Circuits, J. Neurosci., 26, 10380 (2006).

12. A. R. Adamantidis, F. Zhang, A. M. Aravanis, K. Deisseroth, L. de Lecea, Neural substrates of awakening probed with optogenetic control of hypocretin neurons, Nature, 450, 420 (2007).

13. E. Papagiakoumou et al., Scanless two-photon excitation of channelrhodopsin-2, Nat. Methods, 7, 848 (2010).

14. C. W. Freudiger et al., Label-free biomedical imaging with high sensitivity by stimulated Raman scattering microscopy, Science, 322, 1857 (2008).

15. J.-X. Cheng, X. S. Xie, Vibrational spectroscopic imaging of living systems: An emerging platform for biology and medicine, Science, 350, (2015).

16. P. T. So, C. Y. Dong, B. R. Masters, K. M. Berland, Two-photon excitation fluorescence microscopy, Annu. Rev. Biomed. Eng., 2, 399 (2000).

17. D. Fu, G. Holtom, C. Freudiger, X. Zhang, X. S. Xie, Hyperspectral imaging with stimulated Raman scattering by chirped femtosecond lasers, J. Phys. Chem. B, 117, 4634 (2013).

18. D. Fu, X. S. Xie, Reliable cell segmentation based on spectral phasor analysis of hyperspectral stimulated Raman scattering imaging data, Anal. Chem., 86 9, 4115 (2014).

19. J. Shou, Y. Ozeki, Photoswitchable stimulated Raman scattering spectroscopy and microscopy, Opt. Lett., 46, 2176 (2021).

20. F. G. Erko et al., Spectral, Conformational and Photochemical Analyses of Photochromic Dithienylethene: cis-1, 2-Dicyano-1, 2-bis (2, 4, 5-trimethyl-3-thienyl) ethene Revisited, Eur. J. Org. Chem., 2013, 7809 (2013).

21. J. Du, L. Wei, Multicolor Photoactivatable Raman Probes for Subcellular Imaging and Tracking by Cyclopropenone Caging, J. Am. Chem. Soc., (2021).

22. J. Ao et al., Switchable stimulated Raman scattering microscopy with photochromic vibrational probes, Nat. Commun., 12, 1 (2021).

23. S. Habuchi et al., Reversible single-molecule photoswitching in the GFP-like fluorescent protein Dronpa, Proc. Natl. Acad. Sci. U. S. A., 102, 9511 (2005).

24. R. Ando, H. Hama, M. Yamamoto-Hino, H. Mizuno, A. Miyawaki, An optical marker based on the UV-induced green-to-red photoconversion of a fluorescent protein, Proc. Natl. Acad. Sci. U. S. A., 99, 12651 (2002).

25. X. Chen et al., Acetylcholinesterase inhibitors with photoswitchable inhibition of β-amyloid aggregation, ACS Chem. Neurosci., 5, 377 (2014).

26. M. Borowiak et al., Photoswitchable inhibitors of microtubule dynamics optically control mitosis and cell death, Cell, 162, 403 (2015).

27. S. Mondal, S. S. Parelkar, M. Nagar, P. R. Thompson, Photochemical control of protein arginine deiminase (PAD) activity, ACS Chem. Biol., 13, 1057 (2018).

28. B. A. Copits et al., A photoswitchable GPCR-based opsin for presynaptic inhibition, Neuron, 109, 1791 (2021).

