## Supplemental materials for "Real-time precision opto-control of chemical processes in live cells"

The RPOC platform

A femtosecond laser source (InSight X3+, Spectra-Physics) is used for optical signal excitation and opto-control. The laser outputs two femtosecond pulse trains, one at a fixed wavelength of 1045 nm, and the other tunable from 690-1300 nm, both with ~100 fs pulse width. For stimulated Raman scattering (SRS) microscopy, the 1045 nm output is used as the Stokes beam and the tunable output is used as the pump beam. A single 150 mm SF-57 glass rod (Lattice Electro-Optics) is placed in the Stokes beam and two 150 mm SF-57 glass rods are placed after combining the two laser beams for the chirping of the pump and Stokes pulses to 3.4 ps and 1.8 ps, respectively. The laser beams double-pass the two glass rods for additional chirping. An AOM (M1205-P80L-0.5 with 532B-2 driver, Isomet) is used to modulate the Stokes beam for SRS microscopy. Hyperspectral SRS image stack is acquired by tuning the optical delay using a translational stage (X-LSM050A, Zaber Technologies) at 10 µm per step while collecting single-color images. The combined laser beams are directed to a 2D galvo scanner set (GVS002, Thorlabs) and then into an upright microscope (Olympus BX51). Either a 40x/0.8 NA (LUMPLFLN 40XW, Olympus) or a 60x/1.2 NA water immersion objective lens (UPLSAPO 60X, Olympus) is used to focus the laser beams onto the sample. Forward signals are collected using a 1.4 NA oil condenser. A 776 nm long-pass dichroic mirror (FF776-Di01-25x36, Semrock) is used to separate the TPEF signal from the input laser beams. The forward two-photon excitation fluorescence (TPEF) signals and the leaking of the control laser beam are detected after being reflected by the long-pass dichroic mirror. A combination of filters (FF01-575/59-25 or FF01-451/106-25, Semrock; ET425lp, Chroma) is used to detect the fluorescence signal and the leaking of control laser beams. The SRS signals are detected after transmission of the dichroic mirror using a photodiode (S3994, Hamamatsu) paired with a lab-designed tuned amplifier with a center frequency of 2.7 MHz. A short-pass filter (980SP, Chroma Technology) is used to block the Stokes beam from entering the photodiode. A lock-in amplifier (HF2LI, Zurich Instruments) is used to demodulate signals for SRS imaging. The lock-in amplifier and the AOM for Stokes beam modulation are synchronized by a function generator (DG1022Z, Rigol). In the epi-direction, two photomultiplier tubes (PMTs) are installed to collect fluorescence signals at selected wavelength windows using different filters.

A portion of the tunable laser beam and the 1045 nm laser beam is frequency-doubled by BBO crystals (EKSMA Optics) to generate visible wavelengths for opto-control. The crystals are mounted on rotational mounts to optimize the second harmonic generation efficiency at different wavelengths. Flip mirrors are used to select the desired control laser wavelength. The selected visible laser beam is sent to another AOM (M133-aQ80L-1.5 with 522B driver, Isomet) which is controlled by comparator circuit boxes. The optical signal voltage is compared with a preset condition to determine the output TTL voltage for AOM control. The SRS signal output is delivered from the lock-in amplifier, while the fluorescence signal output is delivered from an amplifier (PMT3V4, Advanced Research Instrument Corporation) connected after the PMT.

The design of the comparator circuit box 1 with a single intensity threshold is illustrated in **fig. S1**. Three operation modes of this circuit box include: AOM constantly on, AOM constantly off, and AOM control triggered by the signal-threshold comparison. Threshold voltage $V_T$ can be selected from either a manual tuning knob or a digital input with a range of 0-10V and 0.01 V accuracy. The circuits can be selected from 'AOM constantly on', 'AOM constantly off', and the 'opto-control' modes. The output TTL signal has a <0.7 V output as digital '0' and ~5 V output as digital '1'. The signal output and digital threshold output references are also available.

The design of the comparator circuit with digital logic is illustrated in **fig. S4**. Aside from the same functions as the comparator box 1, a TTL digital input is available allowing this comparator box to be used together with the comparator box 1 for digital logic selections. Digital logic functions can be selected by using jumpers on 3-pin jumper bars. Connections of AND, OR, NAND, and NOR functions are illustrated in **Fig. 3D** and **fig. S10**.

Image acquisition and analysis

Images are saved as .txt files and processed using ImageJ for display. Pseudo-colors are used to represent different chemical compositions for SRS imaging and active pixels (APXs). Spectral or intensity profiles are plotted using Origin Pro. Particle trajectories are tracked using a particle tracker ImageJ plug-in. The parameters to analyze the 100-frame time-lapse SRS image stack and the APX stack are: radius=0, cutoff=3, percentile=0.5, link range=1, displacement=5. A single lipid droplet (LD) trajectory and the corresponding active-pixel trajectory are plotted using ImageJ particle tracker Plug-in together with images for display. Merging different image channels, image subtractions, particle analyses, and intensity integrations are performed using ImageJ built-in functions. Hyperspectral SRS images are analyzed using a spectral phasor plug-in in ImageJ. Plots of chemical maps are pseudocolor-coded for display.

Cell preparation

MIA PaCa-2 pancreatic cancer cells were purchased from ATCC and cultured in Dulbecco's Modified Eagle Medium (DMEM, ATCC) with 10% fetal bovine serum (FBS, ATCC) and 1% penicillin/streptomycin (Thermofisher Scientific). The cells were seeded in glass-bottom dishes (MatTek Life Sciences) with 2 mL culture media and then incubated in a $CO_2$ incubator at 37 °C and 5% $CO_2$ concentration. Cells were grown to about 50% confluency and were directly used for live-cell imaging or fixed with 10% buffered formalin phosphate (Fisher Scientific) for imaging.

Preparation of CMTE and control of CMTE in cells

The chemical *cis*-1,2-dicyano1,2-bis(2,4,5-trimethyl-3-thienyl)ethene (CMTE) was purchased from Sigma Aldrich and prepared in dimethyl sulfoxide (DMSO) at a concentration of 25 mM. MIA PaCa-2 cancer cells were treated with 3.2 µL of the CMTE stock solution for a final concentration of 40 µM. Cells were incubated with CMTE for 8-12 hours before imaging. The combined pump and Stokes laser pulses can gradually switch the CMTE to the closed isomer **1b** with strong signals at 1510 $cm^{-1}$. We deployed RPOC to selectively convert CMTE at different locations of the sample to the open *cis* isomer **1a**, as illustrated in **Fig. 4B**.

ER tracker labeling of cells

MIA PaCa-2 cells were first seeded in glass-bottom dishes and cultured overnight to reach 50%-70% confluency. ER tracker was added to the culture medium with a 3 µM final concentration. The cells were cultured for 30 min at 37 °C and 5% $CO_2$ concentration before imaging. To generate sufficient TPEF signals, femtosecond laser pulses bypassing the chirping rods were directly used for signal generation.

Estimation of the AOM rise time

The AOM rise time satisfies

$$\tau_r = 0.65\frac{d}{V},$$

Here, *d* is the beam diameter, and *V* is the acoustic velocity.

$$d = 2r = 1.22\frac{\lambda}{NA}$$

For the control laser beam at 522 nm and a NA value of 0.01, the beam diameter at the AOM crystal is 0.63 x $10^{-4}$ m. The acoustic velocity (*V*) inside the AOM crystal is 5800 m/s. This gives ~7 ns AOM rise time for the control laser beam.

The design of the comparator circuit box with a single intensity threshold selection

In **fig. S1A**, the optical signal input shown on the left is compared with a voltage threshold that can be set manually using the manual threshold tuning nob or input from the digital threshold input port. The manual & digital switch selects the threshold selection mode. The opto-control TTL signal is output from the right middle port for AOM control. On the right-hand side, two ports are available to deliver the optical signal output and the digital threshold output for references.

The spatial resolution of the imaging system

An SRS image from MIA PaCa-2 cells was acquired to estimate the spatial resolution of the signal generation (**fig. S2**). From the Gaussian fitting of a small feature in the image, the spatial resolution is estimated to be ≤ 373 nm. We estimate the TPEF to have a similar spatial resolution as the SRS.

Beam overlapping and RPOC beam size estimation

Spatial overlapping of the excitation and RPOC laser beams is critical to ensure accurate molecular control. To ensure optimized overlapping between the excitation and RPOC laser beams, we used fluorescence microparticles and compared images using both laser beams. **Fig. S3, A-C** illustrate the condition when the beams are not perfectly overlapped and **fig. S3, D-F** show the condition with optimized beams. Overlap optimization can be achieved by adjusting two mirrors only in the control laser beam path.

From **fig. S3G**, we found that the widths of the same particle imaged using TPEF and 522 nm single-photon RPOC laser are 1.04 and 1.35 µm, respectively. The resolution of the TPEF is determined to be ~373 nm from **fig. S2**. Therefore, the spot size of the RPOC laser ‘x’ satisfies:

$$1.04 = 0.37 \times 2 + a$$

$$1.35 = 2x + a$$

Here ‘x’ is the beam size of the 522 nm RPOC laser beam size, while ‘a’ is the size of the particles excluding the edges. The solution of these equations gives x=525 nm. This beam size is bigger than the theoretical minimum using a NA=1.2 objective lens, majorly due to the reduced beam size of the $1^{st}$ order AOM diffraction as shown in **Fig. 1E**.

The design of the comparator circuit box with the digital logic function

As shown in **fig. S4A**, similar functions to the comparator box 1 with a single threshold selection are available. Besides, a TTL input can be used to perform digital logic calculations with the comparison output from this box. This TTL input can be the TTL output from other comparator boxes. Inverters, an AND gate, and an OR gate are available to achieve different logic combinations.

One function of using two comparator boxes is to select an intensity range for RPOC. The connections of achieving such a function are shown in **fig. S5** and **Fig. 3A**. The upper and lower thresholds are selected by the comparator box 1 and 2, respectively.

The other function of using two comparator boxes is to perform logic calculations from two separate detectors for RPOC. The connections of achieving the AND function are illustrated in **fig. S7** and **Fig. 3D**. Electronic connections for achieving other logic functions are shown in **fig. S10**.

Spectral phasor analysis of cells from hyperspectral SRS images

In **fig. S6**, a single-color SRS image, and segmented images highlighting different organelles including LDs, endoplasmic reticulum (ER), nuclei, cytosol, and a composite image are shown. The ER can also be selected from simply intensity thresholds in SRS images. A single intensity range was used to select APXs on ER in **Fig. 3**.

APXs selection using digital logic signals from two detectors.

A mixture of fluorescent and nonfluorescent polystyrene (PS) microparticles and nicotinamide adenine dinucleotide hydrogen (NADH) crystals is used to demonstrate the digital AND function for APX determination. As shown in **fig. S8A**, an SRS image at the PS aromatic stretching band 3060 $cm^{-1}$ reveals all PS particles in the field of view. In the 450 nm fluorescence channel, both fluorescent PS particles and NADH crystals are visible (**fig. S8B**). The merging of the two channels highlights only the fluorescent PS particles in yellow (**fig. S8C**). Using SRS or fluorescence signals, the APXs can be determined for all PS particles or all fluorescence molecules (**fig. S8, D and E**). Using the AND function, APXs are determined from pixels having signals in both SRS and fluorescence channels, which are the fluorescent PS particles (**fig. S8F**). Adjusting the $V_T$ from two comparator boxes allows optimization of APXs selected using the AND function. **Fig. S9** compares APXs from the AND logic using different intensity thresholds selected for two comparator boxes. The optimal condition for selecting APXs in this case is $V_{T1}$=0.05 V, and $V_{T2}$=0.125 V.

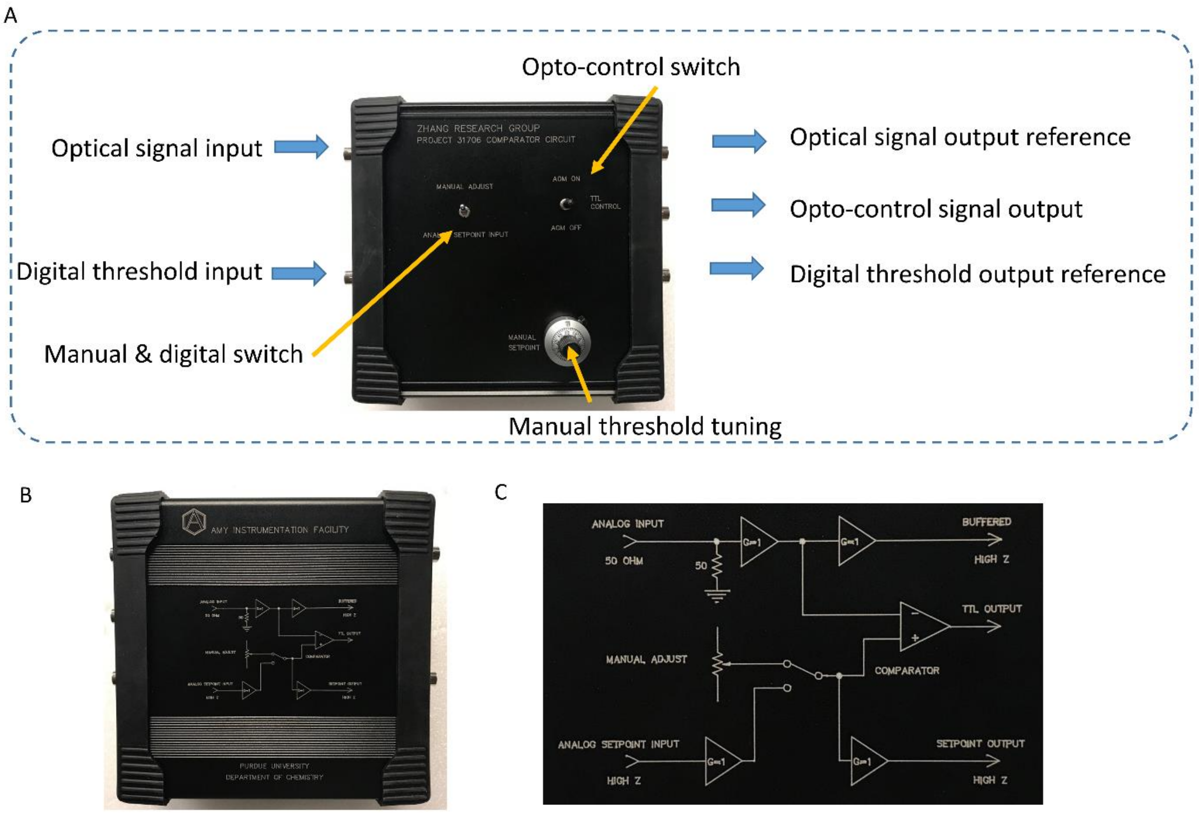


**Fig. S1.** The design of the comparator circuit box 1. (A) The front of the comparator circuit box with explanations of ports and controls. (B) The back of the comparator circuit box. (C) The electronic configuration of the comparator circuit box.

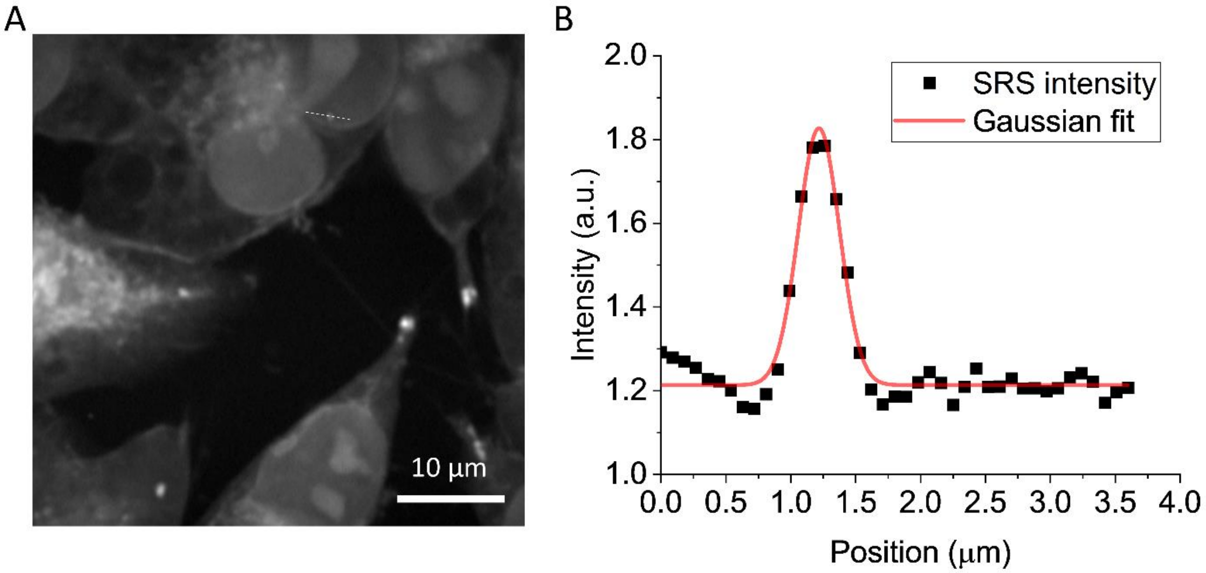


**Fig. S2.** (A) An SRS image of MIA PaCa-2 cells at the 2855 $cm^{-1}$ Raman shift. (B) The SRS intensity profile along the dashed line in panel A. The Gaussian fit quantifies the width of the object, which demonstrates a 373 nm spatial resolution.

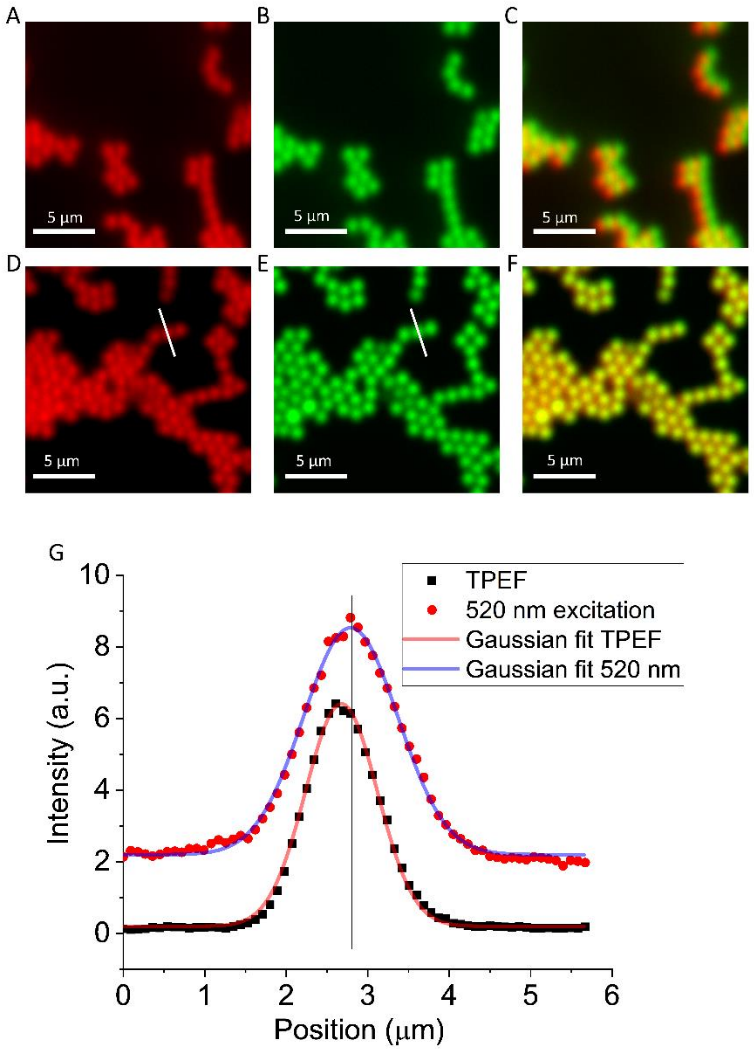


**Fig. S3.** (A) Fluorescence signals from green fluorescent particles excited by the 522 nm RPOC laser beam. (B) TPEF signals from green fluorescent particles excited by the 1045 nm excitation laser beam. (C) Overlay of images in panels A and B, showing an offset due to the misaligned excitation and RPOC laser beams. (D)-(F) Similar images as shown in panels A-C, after optimization of beam overlapping, showing no image offsets in panel F. (G) Single- and two-photon intensity profiles along the lines in panels D and E. The curves are Gaussian fitting results, showing a peak center offset of ~90 nm.

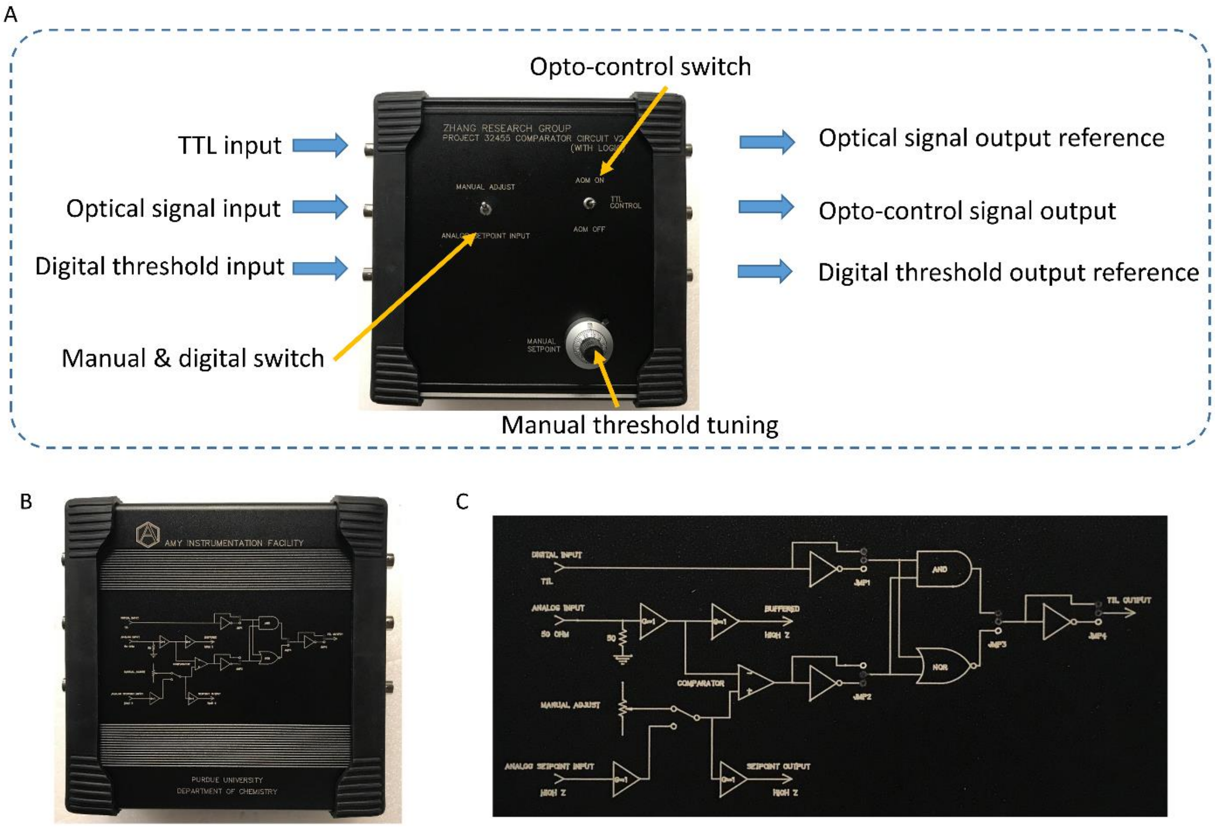


**Fig. S4.** The design of the comparator circuit box 2 with digital logic functions. (A) The front of the comparator circuit box with explanations of ports and controls. (B) The back of the comparator circuit box. (C) The electronic configuration of the comparator circuit box.

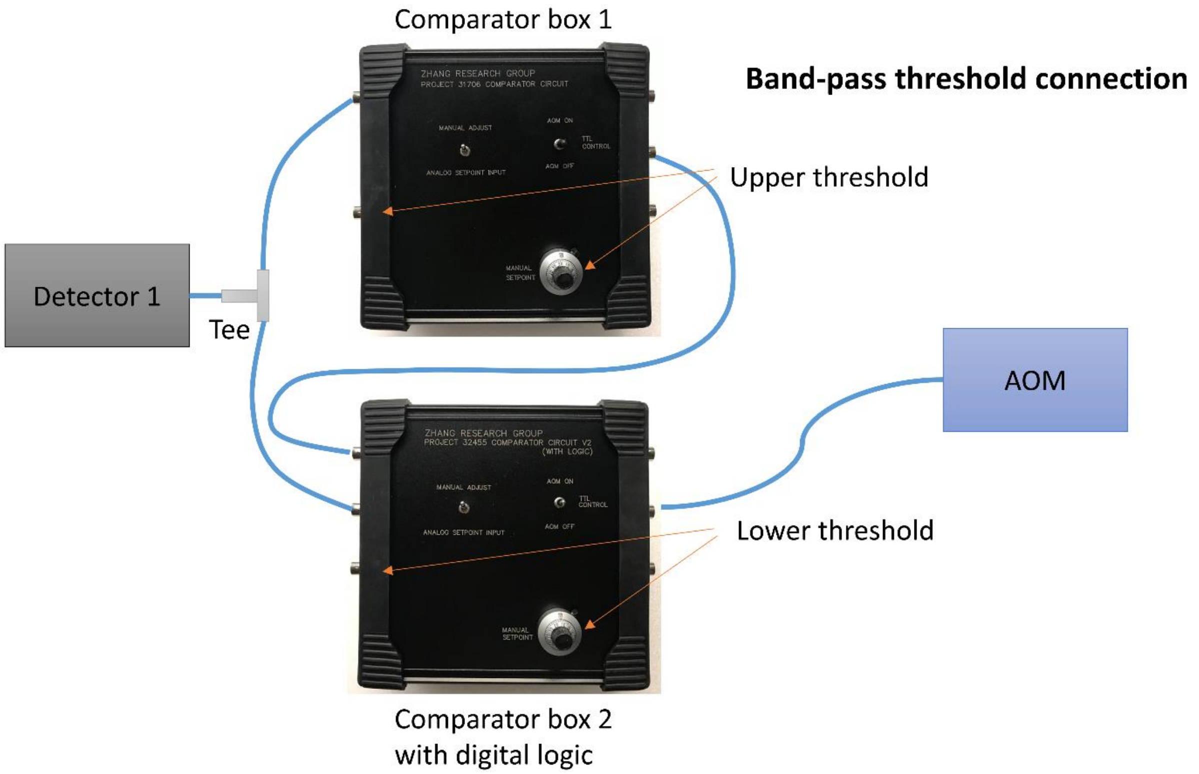


**Fig. S5.** Connections of the two comparator boxes to achieve a band-pass threshold for APX selection.

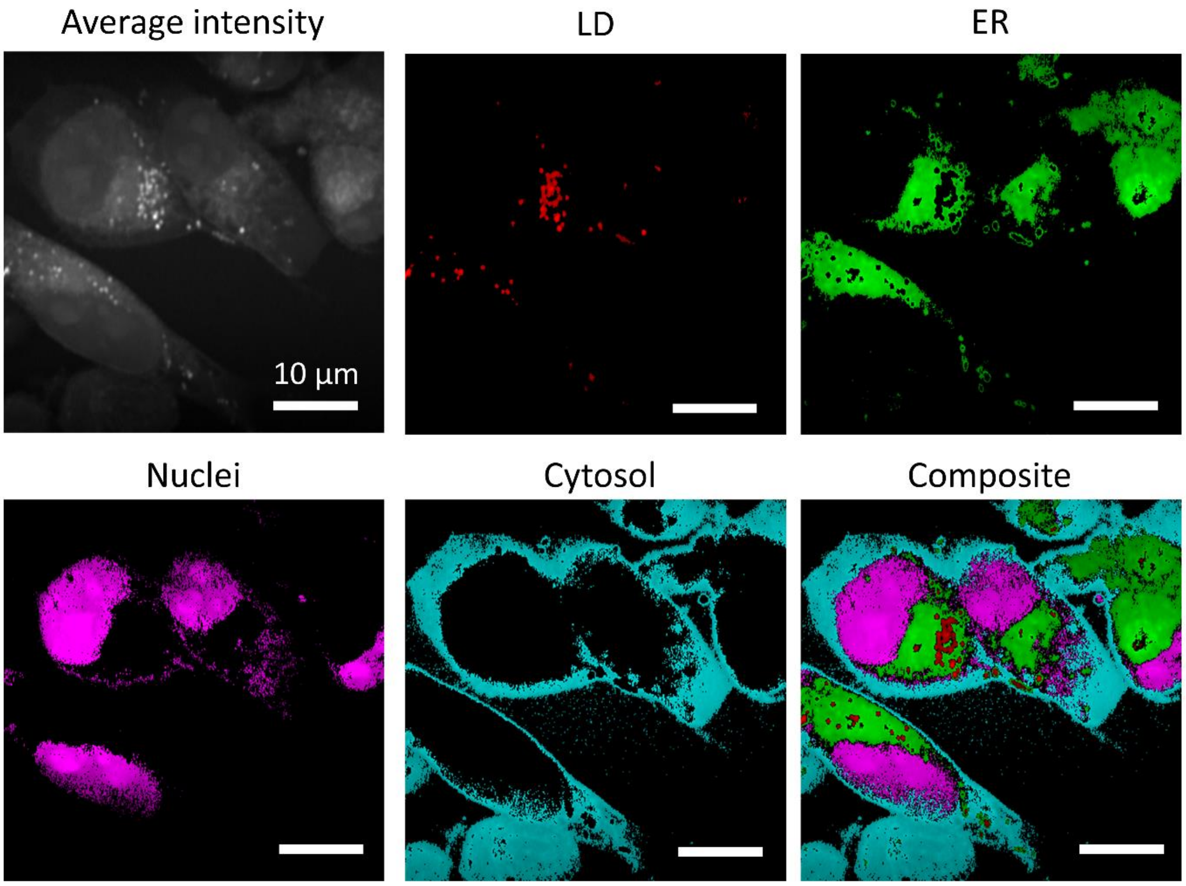


**Fig. S6.** An SRS image (top left) and the chemical maps showing lipid droplets (LDs), endoplasmic reticulum (ER), nuclei, cytosol, and the composite using four chemical compositions generated by spectral phasor analysis of the hyperspectral SRS images.

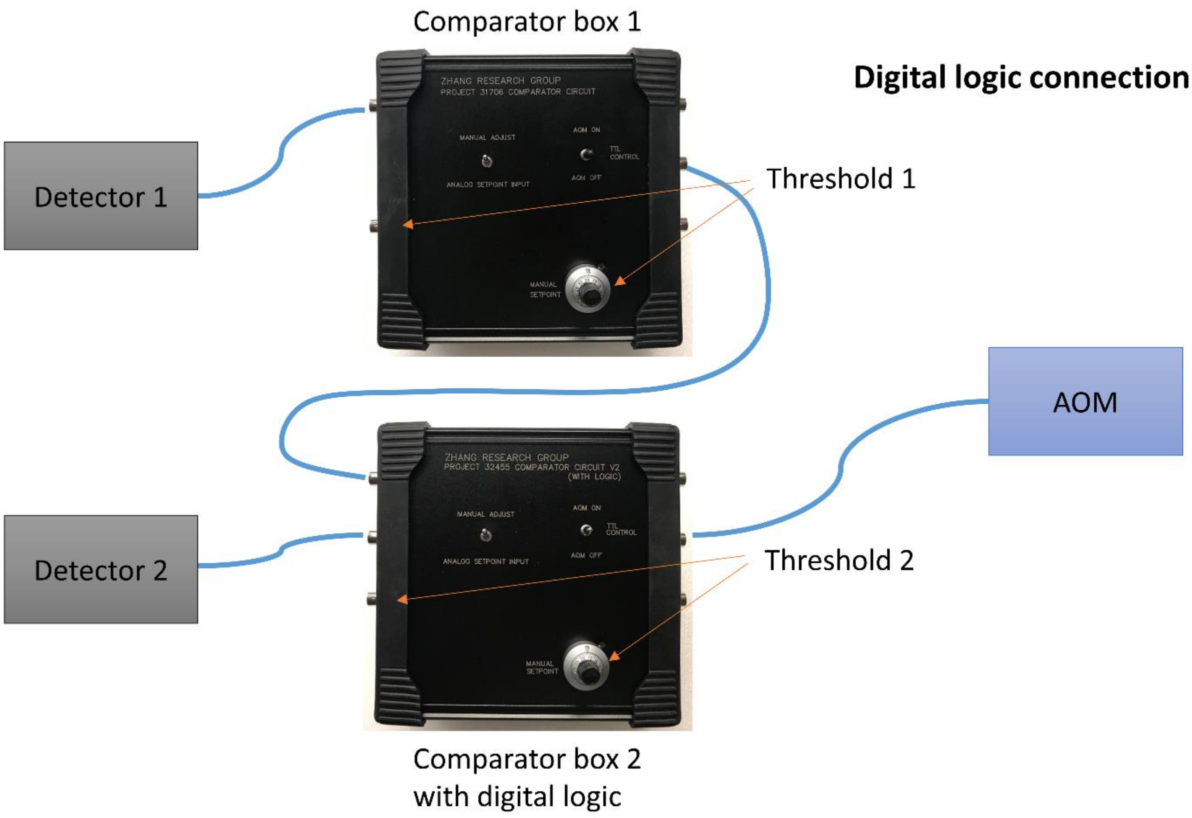


**Fig. S7.** Connections of the two comparator boxes to achieve digital logic functions.

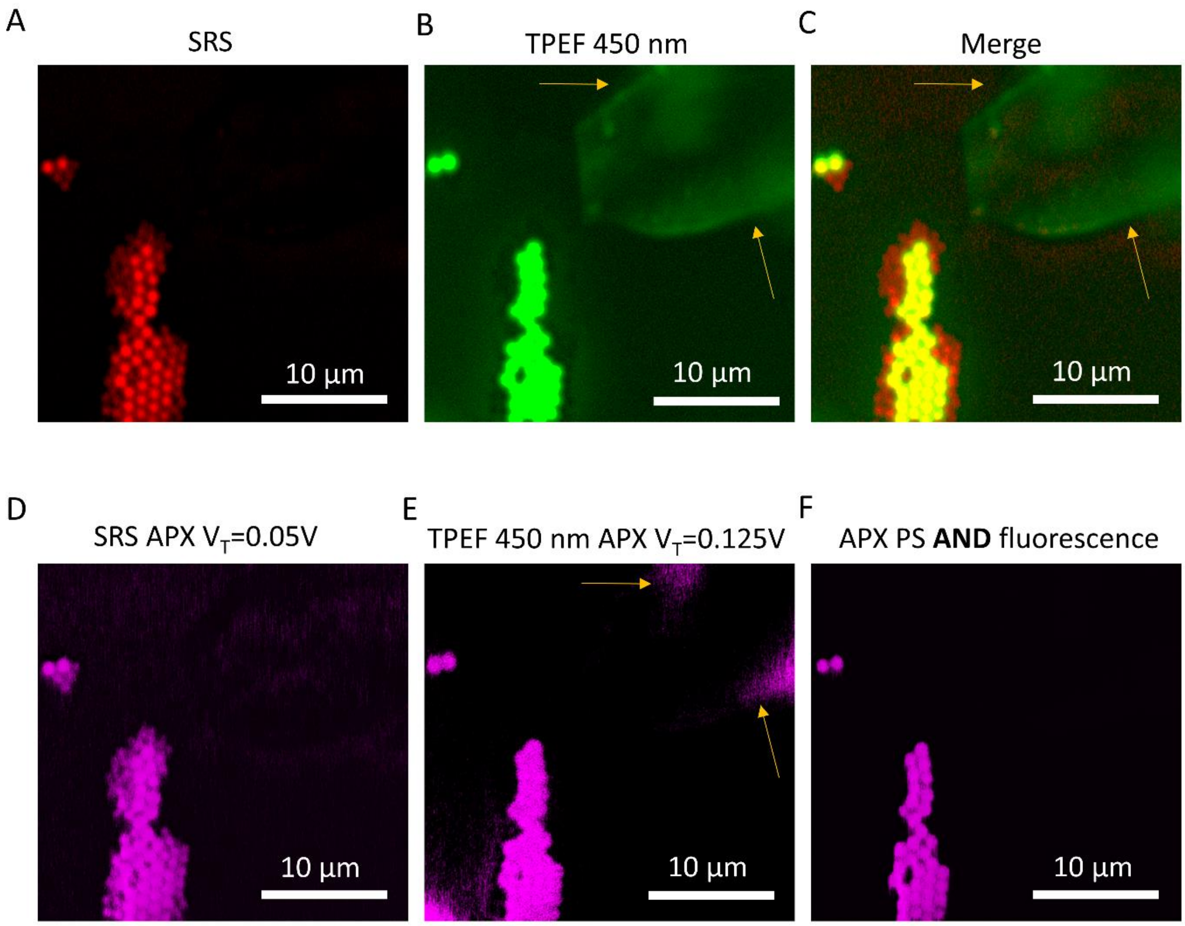


**Fig. S8.** (A) An SRS image of mixed PS particles and NADH crystals at 3060 $cm^{-1}$ Raman shift. (B) A TPEF image from the 450/106 nm channel of the same field of view of the panel A. (C) Merging the SRS and TPEF images from panels A and B. (H) APXs determined by the SRS signals. (D) APXs determined by the TPEF signals. (E) APXs determined by the SRS AND TPEF pixels.

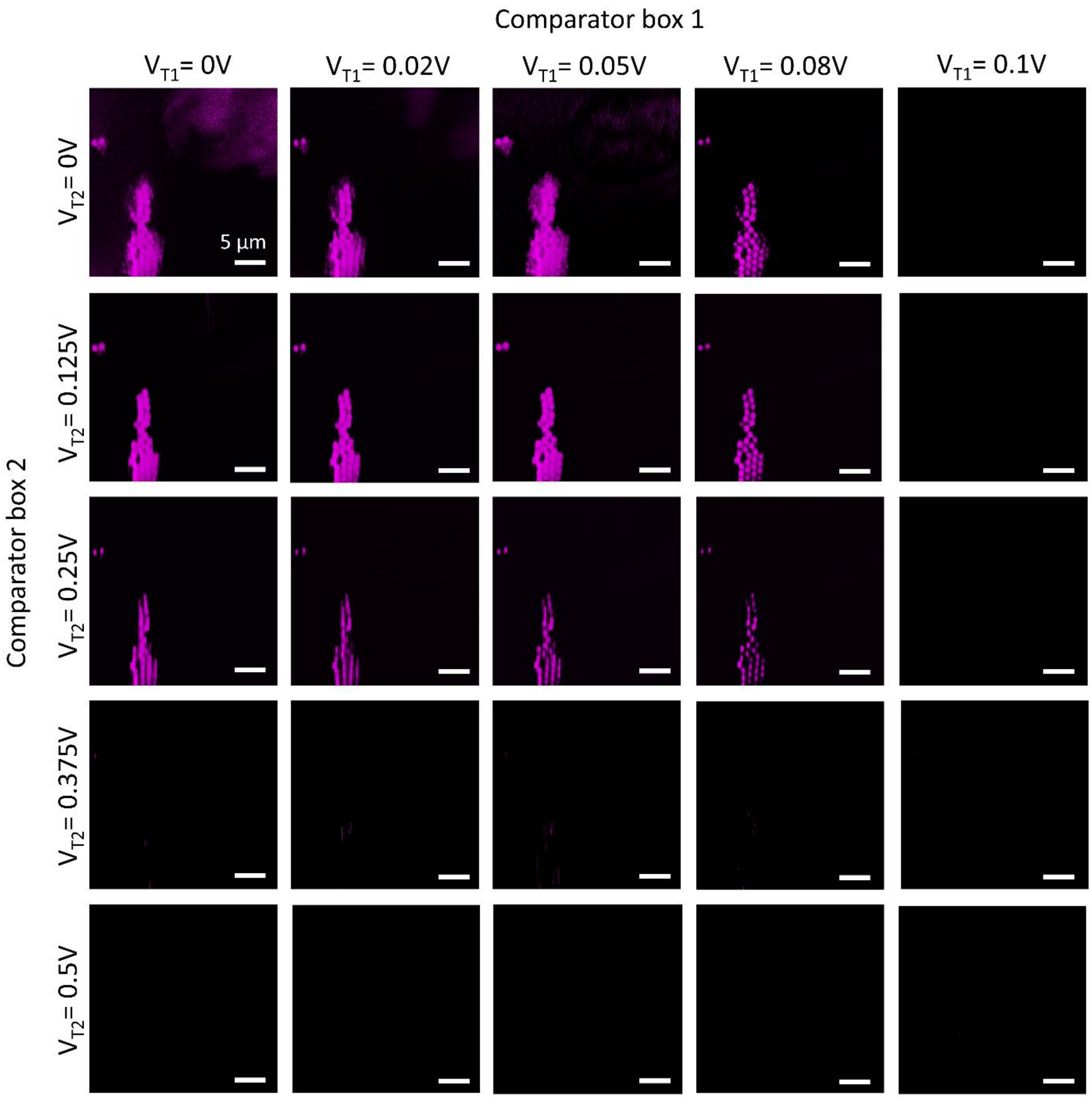


**Fig. S9.** Optimizing the selection conditions using the AND digital function and optimizing the voltage threshold values for each comparator box. The magenta signals are APXs determined in each condition.

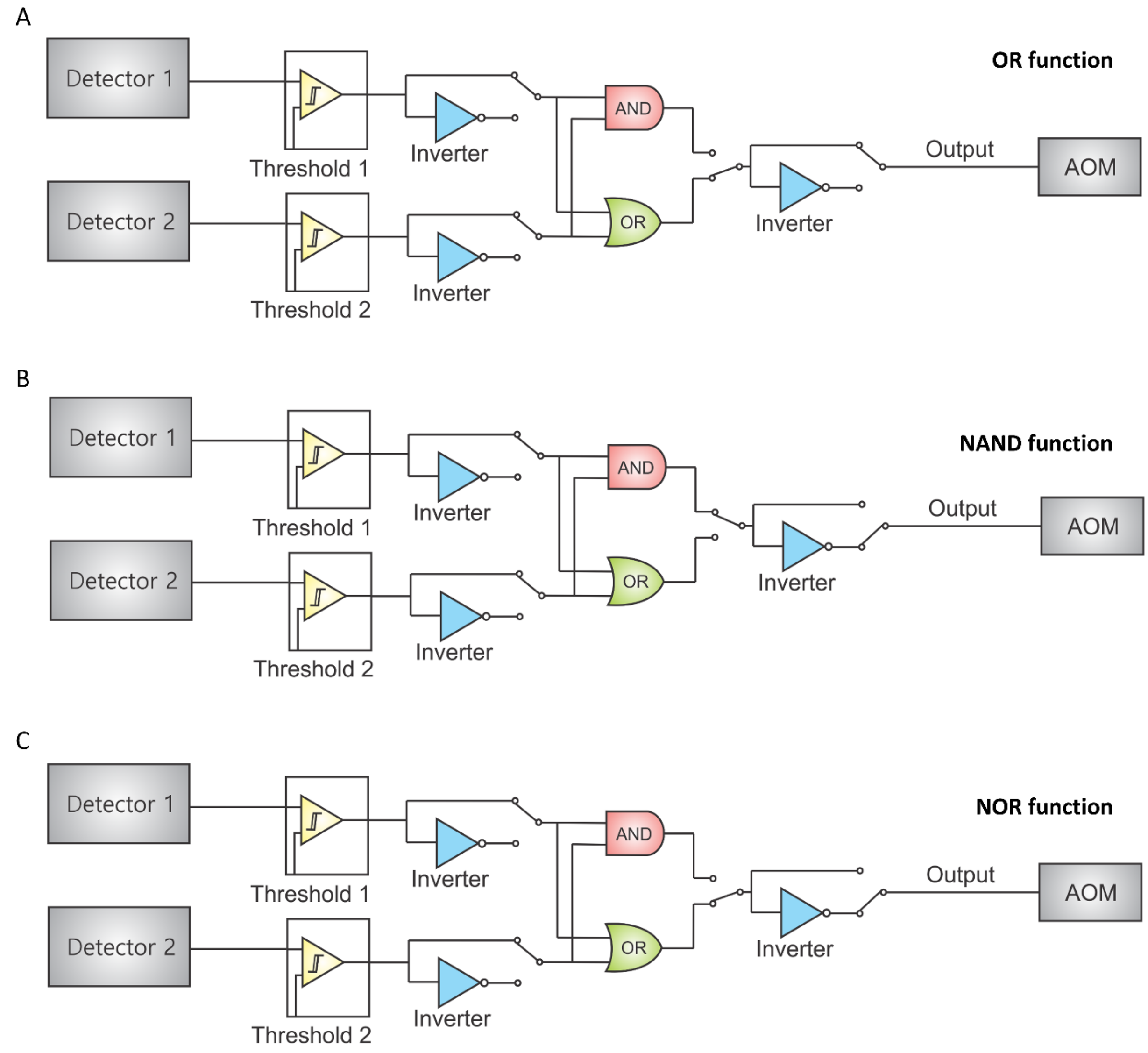


**Fig. S10.** Electronic configurations to achieve the OR, NAND, and NOR logic combinations.

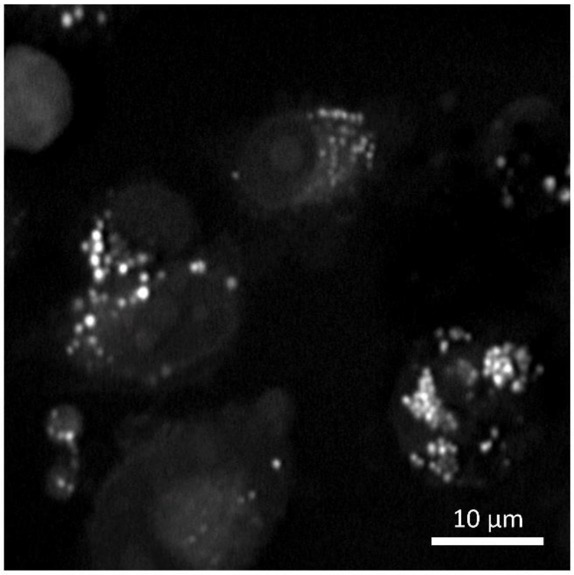


**Fig. S11.** An SRS image of $CH_2$ stretching signals at 2855 $cm^{-1}$ for images in **Fig. 5**.

**Movie S1.** APXs determination by SRS signals from mixed polymer microparticles when scanning the Raman shift from 2800 $cm^{-1}$ to 3100 $cm^{-1}$.

**Movie S2.** Time-lapse SRS images (left) and APXs (right) in live cells at 2.2 s per frame.

**Movie S3.** Time-lapse SRS images (left) and APXs (right) of a single lipid droplet in a live cell at 2.2 s per frame.

**Movie S4.** An illustration of axial scanning for APX determination. Grey: SRS signals at 2855 $cm^{-1}$; Magenta: APXs.

**Movie S5.** An illustration of APXs in a 3D volume. Grey: SRS signals at 2855 $cm^{-1}$; Magenta: APXs.
